# Identity-resolved vocal exchanges in multi-marmoset settings

**DOI:** 10.64898/2026.01.18.699877

**Authors:** J. Cabrera-Moreno, M. Mircheva, J. M. Burkart

## Abstract

Acoustic communication is tightly coupled to collective behaviour and social network structure in many animal societies. Yet continuous, identity-resolved recordings of multiple concurrent signalers are challenging. We evaluated a commercial acoustic camera for marmoset (*Callithrix jacchus*) vocal behavior across controlled bench tests and naturalistic family interactions. Spectral characterization showed the system is closest to neutral and most sensitive in 5–10 kHz, supporting reliable detection without correction in this range. Static localization accuracy was between 1–3 cm across the experimental enclosure. In simultaneous two-source tests, separation depended strongly on geometry and intensity level: very small spacing (20 cm) produced a single merged hotspot, frame-wise co-representation of multiple sources was infrequent and biased by sub-decibel level asymmetries. In natural 15-min family recordings (N=519 calls), the camera detected 96.5% of calls, among detected events, caller assignment was 97.6% correct (overall 94.2%). Ground-truth playback confirmed near-ceiling detection (∼100%) and high assignment with two emitters (∼98%), with reduced attribution in crowded four-emitter layouts. Together, these results indicate that acoustic imaging can deliver high detection and accurate caller attribution for typical marmoset vocalizations, with primary limitations arising from spatial crowding and small inter-source level differences. Importantly, this limitation can be mitigated in the native software by sequentially silencing suspected sound sources. The approach thus provides a practical and reliable path toward continuous, identity-resolved vocal monitoring in freely behaving primates.

## Introduction

In many animal societies, acoustic communication is deeply connected to group behavior and social network structure (Fichtel and Manser 2010). It can facilitate the coordination of group decisions and actions (Briefer et al. 2024), and can play an important role in stabilizing, maintaining, and modifying social relationships (King et al. 2013). Historically framed as dyadic, vocal communication is now understood to unfold in multi-individual settings. Work in group settings shows that, across taxa, third parties adjust signalling to who is present and that bystanders use others’ calls to guide their behaviour, as well as audience and eavesdropping effects (Slocombe and Zuberbühler 2007). Group-level work further demonstrated that vocal signals aid in group cohesion (Gall and Manser 2017), initiate and modulate collective movement (Walker et al. 2017), and coordinate cooperative behaviors like mobbing predators (Gersick et al. 2015). Together, these findings motivate a shift from away from analysing dyadic interactions to a network view of signalling. Recent studies therefore model communication as vocal networks, estimating directed, time-resolved influence among multiple signallers (Stowell, Gill, and Clayton 2016). Yet the evidence remains coarse because caller identity and movement are seldom recorded simultaneously. Moreover, in close-range communication within animal groups, where social interactions naturalistically unfold, it remains challenging to correctly assign a call to the correct caller identity.

To close this gap, two measurement strategies have emerged. Wearable microphones provide individual-level audio in noisy, social settings and can be paired with accelerometers, GPS, or other loggers to enrich behavioural context (Couchoux et al. 2015; Blumstein et al. 2011; Lynch et al. 2013; Demartsev et al. 2024). Their drawbacks include limited battery life, data transfer, potential behavioural impacts, the need to handle animals to fit/remove devices, and limited applicability to small species or individuals that cannot carry tags (e.g., small birds, juveniles). Array-based systems, ranging from custom dense arrays to integrated “acoustic cameras”, offer a fully non-invasive alternative that localizes callers and maps sound sources in space, often with synchronized video (Matsumoto et al. 2022; Guggenberger et al. 2024). Yet, current arrays often lack robust multi-animal identity tracking and rarely report device-level acoustic performance, preventing systematic pairing of calls with movement and transfer for more complex behaviours of vocal exchange and social behaviour. As a result, neither approach has yet standardized nor validated the full technology stack needed to analyse vocal exchanges in naturalistic multi-animal settings.

Integrated acoustic cameras, i.e. packaged microphone arrays with real-time beamforming that overlay acoustic intensity on video, offer a practical instantiation of the array-based approach (Vandendriessche et al. 2021). In principle, this enables non-contact, line-of-sight mapping of who called from where and when, without tags. The approach brings clear strengths, spatialized audio, joint audio-video context, repeatable calibration, but also important constraints: (i) most commercial systems deliver 2D projections (resulting in depth ambiguity when animals align along the camera axis), (ii) usable bandwidth is limited by microphone and sampling-rate choices, (iii) spatial resolution and side-lobe artefacts depend on aperture size and frequency, and (iv) performance can degrade with reverberation and occlusion. Although acoustic cameras have begun to be proposed and used for biological questions (e.g., spatial mapping of callers, non-invasive tracking), systematic validation for multi-animal social vocal scenes is scarce (Guggenberger et al. 2024; Matsumoto et al. 2022; Nagy et al. 2023).

The common marmoset (*Callithrix jacchus*) exemplifies a species with complex behavioral coordination that might be supported by their vocal communication system. Known for their cooperative breeding system (Burkart and Van Schaik 2010; Snowdon and Cronin 2007), high levels of prosociality (Burkart et al. 2007; 2014), and rich vocal repertoire (Grijseels et al. 2023), marmosets engage in complex social behaviours like food sharing, alloparenting, social learning, play, and vigilance (Burkart et al. 2014; French 1996; Burkart, Strasser, and Foglia 2009; Norscia and Palagi 2011; Brügger, Willems, and Burkart 2023), behaviours that are further supported by their sophisticated vocal communication, which includes turn-taking, plasticity in response to noise, volitional control, and labelling of group members (Oren et al. 2024; Roy et al. 2011; Takahashi, Narayanan, and Ghazanfar 2013). Yet much of this work is dyadic and often constrains movement, limiting generalization to natural group interactions.

Here, we provide a targeted, device-level validation of an integrated acoustic camera, an array-based imaging system, for multi-animal vocal scenes in common marmosets. We measure (i) spectral performance with calibrated sweeps and real calls across SPLs (frequency response and sensitivity), (ii) spatial accuracy and beamforming, (iii) source separation across source angles, (iv) localization accuracy in moving emitters, and (v) vocal detection and assignation in real world scenarios during free feeding in a marmoset family group. We report strengths, limits, and best-practice settings for social-vocal experiments, offering a reproducible framework for others.

## Methodology

### Equipment

We used Hextile Nor848B acoustic camera from Norsonic which combines an array of 128 MEMS microphones with imaging software to localise and visually map sound sources in real time. By processing the differences in the arrival time of sound waves at the microphones, the acoustic camera generates a visual representation indicating the exact location of vocalisers. Its frequency range of 20 Hz to 20 kHz encompasses the fundamental frequency and first harmonic of most marmoset vocalisations (Agamaite et al. 2015), while the audio sampling rate of 44.1 kHz ensures high-resolution temporal data. The camera’s wide 105-degree opening angle covers an entire standard experimental enclosure (200 x 100 x 100 cm) from a recording distance of 1 m. With a resolution of 2592 x 1944 pixels and a frame rate of 15 FPS, it captures high-resolution and detailed temporal visual data.

The acoustic camera includes native software for acquisition and review. In this study we relied on five controls used consistently across datasets: (i) Frequency span, which restricts analysis to a chosen band; (ii) Intensity threshold, which filters out any signal below a set level; (iii) Dynamic intensity range, which maps a user-defined dB window to the heat-map palette (e.g., with a 0.5-dB range, a frame peaking at 45.0 dB renders 45.0 dB as red and 44.5 dB as blue); (iv) Virtual microphone, which extracts audio/metrics from a user-placed point; and (v) Suppression point, which masks a selected location to suppress the visualisation of a source. Specific settings and their use are detailed in the Data Collection and Analysis sections.

As a reference microphone we used an Avisoft-Bioacoustics CM16/CMPA. Two monitor speakers (KRK Rokit 5) were used to deliver the acoustic stimuli.

### Acoustic stimuli

The sinewave (1 kHz at -20dB FS), exponential (logarithmic) sine-sweep (1–30 kHz of 10 sec), and band-limited noise bursts (1 – 30 kHz of durations: 100ms, 250ms and 500ms) were synthesized using a custom-made script on Python, relying on the libraries Numpy, Soundfile and Wave.

The marmoset vocalizations used were recorded as part of a different lab’s study on marmoset volubility in different contexts. These recordings were conducted in a separate experimental room where animals were led in pairs. All occurring vocalizations from both individuals were recorded in multiple 20-minute test sessions using two AviSoft Bioacoustics CM16 microphones positioned 1.5m from the experimental enclosure, each connected to an AviSoft Ultrasound Gate 116 at a sampling rate of 192 kHz. Recordings were processed with AviSoft-SAS-Lab Pro (Version 5.3.01) by manually assigning call types based on the time-frequency representations of the vocalizations.

### Animals

A family of 4 adult common marmosets (*Callithrix jacchus)* of either sex was involved for the *Real-world vocal dynamics* test. Animals were housed inside an enclosure of 270 x 180 x 240 cm, with access to an outer enclosure of 320 x 180 x 240 cm when weather conditions allowed. Neighbouring families were housed in adjacent enclosures on either side, without visual contact. The animals were fed mash in the morning and a variety of fresh and cooked vegetables at midday, followed by a snack (e.g., egg, crickets, nuts, or cheese), Arabic gum, and mealworms in the afternoon. Water was provided ad libitum. All animal procedures of this study were approved by the responsible regional government office under the license number 35517.

### Experimental setup

All the acoustic tests took place inside an experimental grid-enclosure placed inside an acoustic chamber (350 x 340 x 230 cm). For the experiments with the marmoset family, animals could enter the experimental grid-enclosure (200 x 100 x 100 cm) through a tubular system connecting their home enclosure with the acoustic chamber and the experimental grid-enclosure. Animals remained in the experimental enclosure for 15 minutes. Mealworms and solid pieces of Arabic gum were provided *ad libitum* to the animals during the testing sessions.

### System calibration

We calibrated the dB SPL delivered by the speaker using a sine wave of 1 kHz at - 20dB FS. The speaker was positioned 1m away from the SPL meter at a similar height at 1.10m and an intensity of 85 dB SPL was measured. Once this intensity was reached, the position of the speaker was maintained for future measurements, as well as the volume settings of the MacBook Pro in charge of running the Python code for the playbacks. The position of the SPL meter was replaced by the acoustic camera. The reference microphone was placed 30 cm away from the center to the left of the acoustic camera. To vary dB SPL, we used the speaker’s manual gain knob in 0.1 dB steps from +10 dB to −70 dB. Because the knob is an analog gain control, every 1 dB turn changes acoustic SPL by exactly 1 dB. Importantly, the dB SPL reported by the acoustic camera matched the intensity measured by the SPL meter, indicating that the camera software retrieves SPL values consistent with the external meter.

### Validation tests

To validate the performance of the acoustic camera in capturing and localizing the vocalizations of multiple marmosets we ran a series of tests assessing for 1) Frequency response and sensitivity, 2) Spatial localization accuracy, 3) Source separation of static, simultaneous emitters, 4) Dynamic source separation of moving emitters, and 5) Real-world marmoset vocal dynamics.

#### 1. Frequency response and sensitivity

We assessed the acoustic camera within the marmoset band using two stimulus classes, an exponential sweep (1–30 kHz, 10 s) and pre-recorded vocalizations (three exemplars each of phee, twitter, food, and tsik-ek vocalisations, 12 total). Each stimulus was presented at 80-, 60-, 40-, and 20-dB SPL, with five repetitions per level. The playback, the reference microphone, and the acoustic camera shared the same geometry as during calibration.

Because the reference microphone has reduced low-frequency sensitivity, all analyses were a priori limited to 2–20 kHz. Audio was sampled at 192 kHz and power spectral densities (PSD) were estimated with Welch’s method (Hann, nfft = 4096, 50% overlap). For each device 𝑑, frequency response deviation (FRD) was defined as

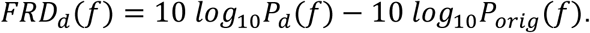

To isolate the camera from room/loudspeaker effects we used the differential deviation

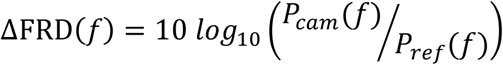

and reported 1/3-octave–smoothed curves. For summary metrics we computed root mean square (RMS) ΔFRD and max ∣Δ∣ within 2–5, 5–10, 10–15, 15–20 kHz bands and 2–20 kHz overall, then took medians across stimuli.

For sensitivity, we recorded a camera background/noise track to obtain 𝑃_𝑛𝑜𝑖𝑠𝑒_(𝑓). For each stimulus level we formed SNR in dB as 𝑃_𝑐𝑎𝑚_(𝑓) − 𝑃_𝑛𝑜𝑖𝑠𝑒_ (𝑓) and interpolated the minimum detectable SPL at SNR = 6 dB. Thresholds were summarized by the same frequency bands using the median across stimuli. To check stimulus dependence, we compared bandwise neutrality metrics between the sweep and the pooled vocalizations.

For the frequency response sensitivity as well as for the remaining tests, fixed acquisition parameters were set in the native acoustic camera software at the beginning of each test. (i) Frequency span: 1 kHz – 20 kHz; (ii) Intensity threshold: 35 dB SPL; (iii) Dynamic intensity range: 0.5 dB – 1.0 dB; (iv) Virtual microphone: at the center of the emitter or in the middle of both; and (v) Suppression point: only used in the natural setting and the ground truth.

#### 2. Spatial localization accuracy

With the operative band established, we next asked how precisely the system localizes within the experimental enclosure, since identity resolution ultimately depends on spatial precision across positions and call types. For this test, we assessed 2D localization accuracy by comparing the acoustic camera’s heatmap maximum to a known loudspeaker position in the image plane. For each of five fixed speaker locations within the enclosure (upper-left, upper-right, center, lower-left, lower-right), we presented 250-ms band-limited noise bursts (5–30 kHz, 70 dB SPL) and five randomly selected exemplars per call type (phee, twitter, food, tsik-ek; 70 dB SPL), each repeated five times, while screen-capturing the camera’s display. On the first frame of each recording, we manually delimited a region of interest (ROI) around the loudspeaker and marked the tweeter center and a point on its rim, yielding both the ground-truth center in full-frame pixels and the tweeter diameter in pixels for metric conversion.

Heatmaps were segmented in Hue, Saturation, and Value (HSV) color space using interactively tuned thresholds optimized on the ROI, then applied frame-by-frame to the full recording. We retained the largest connected hot region per frame and computed its centroid and area in pixels, mapped back to full-frame coordinates by the ROI offset. Localization error for each frame was defined as the Euclidean distance between that centroid and the pre-marked tweeter center. Pixel distances were converted to centimeters using a single scale factor derived from the known tweeter diameter (cm) divided by its measured diameter (pixels), assuming negligible perspective distortion within the ROI. The same procedure was executed independently for each of the five speaker positions, and framewise distances were subsequently summarized per stimulus and position to yield localization accuracy in centimeters.

#### 3. Source separation of static, simultaneous emitters

To assess the ability of the acoustic camera to differentiate simultaneous vocalizations from multiple sources we ran the following tests:

##### 3.1 Simultaneous emission at different distances

To test the camera’s ability to separate two concurrent sources, we presented simultaneous burst trains from two loudspeakers positioned inside the experimental enclosure at fixed, well-separated locations (20, 40, 60 and 80 cm; height matched to the camera’s midline). Both speakers were driven with independent band-limited noise (identical spectrum, different random seeds) to avoid coherence artefacts. Each train comprised five 500-ms bursts separated by 100-ms gaps, presented at 70 dB SPL per speaker (SPL calibrated at 1 m as in the intensity calibration section). We recorded N=5 trains per session.

During each train we saved the camera’s intensity heatmaps/videos and processed them with the same fixed pipeline used in the *Spatial location accuracy* test: a hand-defined ROI around the speakers, HSV thresholding (thresholds tuned once on a held-out frame and kept fixed across all runs), and computation of blob centroids area and distance to true center per frame. We retained up to the two largest blobs per frame and converted their centroids and distance to centimetres.

For every frame in which both bursts were “on”, we attempted to assign two blobs to the two speakers. A blob was considered a correct detection for a given speaker if its centroid lay within the ROI of that speaker’s ground-truth position (radius chosen a priori from the peripheral localization error observed in *Spatial location accuracy*). Frames were classified as: (i) two-peak success (both speakers detected), (ii) miss (one or both speakers absent), or (iii) swap (a single blob closer to the wrong speaker or two blobs both assigned to the same speaker).

The primary endpoint was the two-peak detection rate (percentage of “on” frames with two-peak success). Secondary endpoints were per-speaker miss rate, swap rate, and localization error distribution during simultaneity (median and IQR per train). To link confidence to performance, we computed the correlation between blob area and localization error across frames.

##### 3.2 Level asymmetry

Using the same noise burst-trains as for *Simultaneous emission at different distances,* we tested next for inter-source level differences. One “reference” speaker was held at 70 dB SPL, the other (“test”) was attenuated by −0.2, −0.5, and −1.0 dB SPL relative to the reference. The distance between the speakers was fixed at 70 cm with the similar height as for the calibration procedure. All acquisition steps matched *Simultaneous emission at different distances*, including ROI definition, centroiding, pixel-to-cm scaling, and framewise outcome extraction. Offsets were randomized across runs, with ≥5 trains per offset. For each frame we scored per-speaker detection (hotspot within the pre-registered ROI) and two-source detection (both ROIs detected), and computed localization accuracy as centroid-to-speaker distance (cm) on detected frames. Primary outcomes were per-speaker detection rates, the two-source fraction, and localization-error distributions (median, IQR) for each offset.

#### 4. Dynamic source separation of moving emitters

Because callers may not always be static, we then tested whether localization remains stable as a source translates, probing whether the static accuracies generalize to motion. Here, we evaluated the camera’s ability to track a moving source and localize it over time. A loudspeaker (as in previous tests) was hand-moved along a planar path covering the full enclosure length (left to right and right to left) while emitting band-limited noise trains at 70 dB SPL (five 500-ms bursts separated by 100-ms gaps). Because motion was not scripted, kinematics was estimated offline: for each burst (“bout”), we took the first and last frames with valid data to compute displacement (centroid position*_last_* – centroid position*_first_*) and duration (time*_last_* – time*_first_*), yielding a bout-wise speed. Pixel-to-centimeter conversion used the enclosure grid as a physical ruler.

Acquisition settings (dynamic frequency span, intensity range/thresholds) were followed by 2 or 3 tests. Unlike static tests, the emitter’s image location varied, we therefore used a YOLOv8 detector–tracker to locate the loudspeaker in each frame and derive its geometric center. Detector performance was validated against 200 manually annotated frames, achieving 99.1% detection with a maximum center deviation of 0.2 cm relative to the annotations. After center estimation, processing followed the same procedure as in test *Spatial location accuracy,* where for every frame we extracted the acoustic heatmap, applied the same HSV threshold, computed the centroid of the largest blob and calculate the Euclidian distance to the center in cm. Each noise bout was treated as an independent observational unit, carrying its own speed and localization accuracy, all frames were indexed to their bout ID to permit within-bout analyses of error and detection continuity.

#### 5. Real-world marmoset vocal dynamics

Finally, we assessed end-to-end performance in a freely moving marmoset family (N=4; breeding pair + two helpers) during a 15-min free feeding session, using the same camera placement, array settings as in prior tests. Animals accessed the experimental enclosure via the routine tube system used in their daily procedures. Two feeding stations (front-upper, back-lower) encouraged depth separation to reduce occlusion.

Audio–video acquisition used the camera’s native recording. Raw audio was exported as WAV and segmented with WhisperSegg (Gu et al. 2023) (pretrained for marmoset calls, default operating point, ∼93% reported accuracy) to obtain time-stamped call candidates and automatic call-type labels. Segmentation outputs (CSV) and the original WAV were imported into ELAN together with the camera video. Through manual annotation we (i) verified call types and (ii) assigned a caller identity (animal ID) when the heat-map focus unambiguously coincided with a visible animal. Each event was given a confidence label: certain (a single, well-defined hotspot on a single animal) or uncertain (ambiguous hotspot position or no hotspot with only a spectrographic trace).

Primary outcomes were: (1) detection rate = proportion of calls with a corresponding camera hotspot (spatially around an animal), (2) assignment accuracy = proportion of detected events correctly attributed to the manually verified caller; and (3) per-type performance (precision/recall/F1 for phee, twitter, food, tsik-ek).

To avoid bias, no post-hoc retuning of camera parameters was used for the primary analysis. In ambiguous cases flagged during adjudication, we allowed a secondary adjudication pass in the native software (adjusted frequency span, intensity threshold, dynamic range, virtual microphone, suppression point, and array distance) solely to classify those events as probable or uncertain.

##### Controlled playback for ground truth

To benchmark assignment under known identities, we created four 5-min playback streams comprising natural calls (food, phee, tsik-ek) from five individually recorded animals. Streams were level-matched (A-weighted 70 dB SPL at 1 m) and mapped to distinct loudspeakers, so each speaker corresponded to a fixed “animal ID.” We recorded two scenarios: (i) two-speaker layouts at four positions (front-close ∼5 cm apart; front-spaced ∼20 cm; back-close; back-spaced), and (ii) four-speaker layouts replicating the spatial distribution observed during free-feeding. Layout order was not randomized and followed the following sequence: front close, front far, back far, back close, transitions were logged with absolute timestamps. Analysis mirrored the naturalistic pipeline: a detection was a hotspot around the active speaker during each call window, assignment was correct when the hotspot matched the emitting speaker (5cm radius). We summarized detection and assignment rates overall and by layout, with bootstrapped 95% CIs, and mixed-effects models (random intercepts for speaker and call exemplar) were used to compare layouts while accounting for repeated calls per source.

## Results

### 1. Frequency response and sensitivity

The camera is most spectrally neutral in the 5–10 kHz band, where its response shows the least coloration (Figure 1 C). Across all stimuli, the typical deviation from neutral (RMS ΔFRD) over 2–20 kHz was ∼6.2 dB, but within 5–10 kHz it dropped to ∼2.3 dB (worst case 3.9 dB). Outside this band, deviations increased: ∼4.9 dB in 2–5 kHz (max 6.0), ∼6.8 dB in 10–15 kHz (max 8.4), and ∼8.6 dB in 15–20 kHz (max 8.7). Thus, 5–10 kHz is the flattest and most reliable operating range.

**Figure 1.**
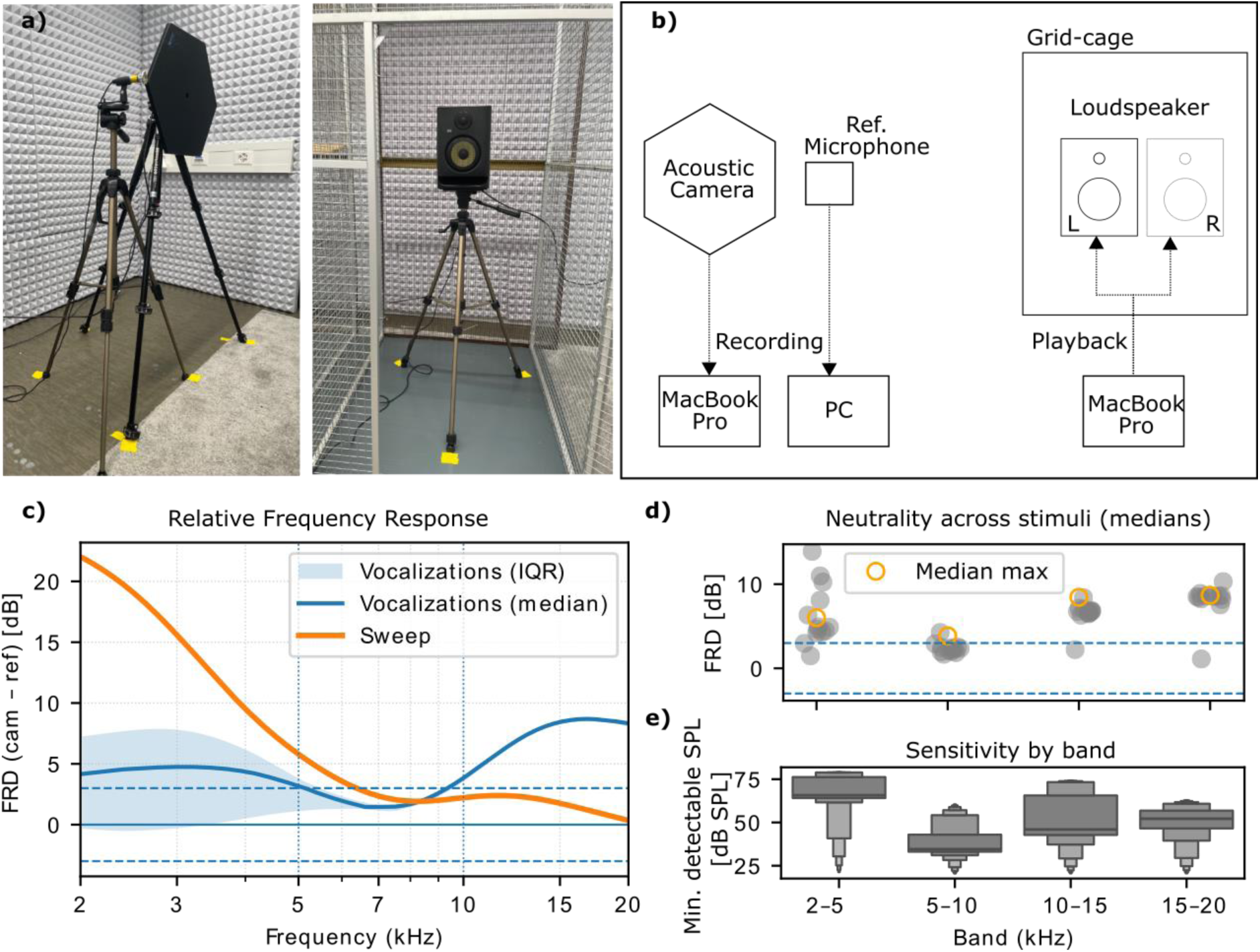
a) Placement geometry of the acoustic camera together with the reference microphone and the playback speaker inside the acoustic chamber. b) Setup connectivity. c) Relative frequency response (ΔFRD) of the camera vs the reference mic (2–20 kHz, 80 dB SPL, 1/3-oct). Thick lines: sweep and median vocalizations, shaded band: IQR across vocalizations, dashed: ±3 dB. d) Bandwise neutrality across stimuli: median RMS ΔFRD (grey points), hollow points show median max |Δ|. e) Sensitivity by band: median minimum detectable SPL (SNR = 6 dB).

We also tested whether these neutrality estimates depended on the stimulus. In 5–10 kHz, sweep-based and pooled natural-call estimates were statistically indistinguishable (Kruskal–Wallis p = 0.77). Differences were larger outside 5–10 kHz, but when averaged over 2–20 kHz, the two methods remained closely aligned (median difference 0.22 dB) (Figure 1 C). Although overall curve shapes differed (Pearson r ≈ −0.28), the practical conclusion is stable: performance is best in 5–10 kHz and degrades toward the band edges.

Sensitivity showed the same pattern. Minimum detectable levels at SNR = 6 dB (median across stimuli) were ∼65.7 dB SPL in 2–5 kHz, ∼34.5 dB SPL in 5–10 kHz, ∼45.9 dB SPL in 10–15 kHz, and ∼52.2 dB SPL in 15–20 kHz (Figure 1 D, E). The lowest thresholds in 5–10 kHz indicate maximal sensitivity where marmoset calls carry most energy, with progressively higher thresholds at the band edges.

### 2. Spatial localization accuracy

Using event-level localization error (median distance from the detected hotspot to the loudspeaker), accuracy depended strongly on position in the enclosure and, to a lesser extent, on call type. A two-way model (location × call type) showed large main effects of location (F(4,300)=936.6, p<10⁻¹⁶⁰) and call type (F(4,300)=57.4, p≈6.5×10⁻³⁶), with a significant interaction (F(16,300)=33.0, p≈1.4×10⁻⁵⁶). Deviations were minimal at the center (mean±SD 0.27±0.09 cm; median 0.25 cm) and increased toward the corners/edges (down-left 2.40±0.16 cm; down-right 2.29±0.29 cm; up-left 2.68±0.40 cm; up-right 2.31±0.90 cm).

Across call types (pooled over locations), phee calls were localized more accurately tsik-ek calls (mean difference ≈0.56 cm, Tukey p=0.004). Other pairwise contrasts were not significant.

Overall, the camera localizes sources with sub-centimeter accuracy at the enclosure center and ≈2–3 cm near the periphery, performance varies modestly with call type, with phee yielding the smallest errors.

### 3. Source separation of static, simultaneous emitters

#### 3.1 Simultaneous emission at different distances

We presented two synchronous noise-burst trains through spatially separated speakers and quantified, per video frame, whether the acoustic camera produced a detection for the left and/or right source and how far the detected hotspot lay from each speaker.

At 80 cm separation, the camera detected the left and right sources in 50.95% and 57.66% of frames, respectively (Wilson 95% CI: 46.89–54.99% and 53.60–61.61%), two-source frames occurred in 10.15% (8.0–12.9%). When detected, the median distance from hotspot to the true speaker was 2.50 cm (IQR 0.07) on the left and 3.63 cm (IQR 0.08) on the right, indicating accurate localization on detected frames.

At 60 cm, detections were asymmetric: left in 88.98% (86.18–91.28%) of frames and right in 33.91% (30.17–37.85%), two-source detections appeared in 24.44% (21.12–28.10%). Median distances on detected frames were 2.04 cm (IQR 0.07; left) and 2.75 cm (IQR 0.67; right).

At 40 cm, the asymmetry flipped: right was detected in 91.22% (88.64–93.26%) of frames and left in 40.96% (37.0–45.0%), two-source detections increased to 33.2% (29.5–37.1%). Median distances on detected frames were 1.61 cm (IQR 0.37; right) and 1.87 cm (IQR 0.06; left).

Finally, at 20 cm, the camera detected activity for both sides in essentially all frames (two-source: 100.0%, 99.34–100.00%; left: 100.0%, 99.34–100.00%; right: 100.0%, 99.34–100.00%). However, inspection of x/y centroids showed that the two heatmap blobs collapsed to a single hotspot midway between the speakers (identical centroids across sides), so distances to the physical speakers were larger despite robust detection (median 10.95 cm, IQR 0.59; left) and (8.18 cm, IQR 0.62; right). We therefore interpret 20 cm as a merged-source regime: the camera is highly responsive but does not resolve two distinct, simultaneous peaks. In practice, this ambiguity can be mitigated using the acoustic camera’s native software by placing a “silence point” over one source, which disambiguates the other. However, to maintain a uniform analysis across bench tests, we did not apply this post-hoc retuning and used it only in the naturalistic sessions.

In summary, across spacings, the system localizes detected sources with centimeter-level precision, while the fraction of frames with simultaneous two-source detections decreases from 100% (merged hotspot at 20 cm) to 33% (40 cm), 24% (60 cm), and 10% (80 cm). This pattern is consistent with array-resolution limits: very small spacing produces a single merged hotspot, moderate spacing yields frequent single-source detections with a minority of frames showing two distinct peaks, and wide spacing reduces the coincidence of two detections in the same frame, even though localization remains accurate when a blob is present.

#### 3.2 Level asymmetry

With two simultaneous noise-burst trains and fixed geometry, even a 0.2 dB inter-speaker level offset sharply reduced two-source detections (0.9% of frames; 95% CI 0.4–2.0%) and biased detections toward the slightly louder source (left 94.5% vs. right 6.3%). At 0.5–1.0 dB offset, two-source frames were essentially absent (0.0% [0.0–0.7%]), with the louder source detected on 98.6–99.3% of frames and the attenuated source on 0.7–1.4%. Localization error for the detected louder source remained small (median ≈2.6 cm; narrow IQR), whereas the attenuated source showed larger and less stable errors owing to sparse detections. These asymmetries reflect a winner-takes-most map rather than faulty centroids. Note that manual parameter retuning in the native software can expose the weaker source.

### 4. Dynamic source separation of moving emitters

We analyzed 50 motion bouts while the speaker traversed the enclosure emitting five 500-ms noise bursts (100-ms gaps). Bout speeds spanned 0.06–21.21 cm/s (median = 3.88 cm/s, IQR = 10.76 cm/s). Localization accuracy, indexed as the median distance between the heat-map centroid and the speaker’s center per bout, was 2.53 cm (IQR = 0.90 cm; 95th percentile = 3.56 cm). Across bouts, accuracy did not degrade with speed: Pearson r = 0.137 (p = 0.343) and Spearman ρ = 0.009 (p = 0.951) between speed and median error. A simple linear model estimated a slope of 0.0137 cm error per 1 cm/s (95% CI: −0.015 to 0.042 cm/s; p = 0.343), i.e., no statistically reliable dependence. Overall, the camera maintained ∼2.5–3.5 cm localization error over the observed speed range, indicating stable tracking performance during naturalistic, non-constant motion.

### 5. Real-world marmoset vocal dynamics

Across the 15-min family recording (N = 519 calls identified in the spectrogram), the acoustic camera flagged a spatial hotspot for 501 calls, yielding a detection rate of 96.5% (Wilson 95% CI: 94.5–97.8%). Among detected calls, assignment accuracy, the hotspot landing on the manually verified caller, was 97.6% (489/501; 95% CI: 95.8–98.6%). (Overall correct assignments relative to all calls were 489/519 = 94.2%.).

By call type, detection and assignment were for Food calls (fc, n=224): detection 97.8%, assignment 98.2%. Phees (ph, n=155): detection 98.1%, assignment 99.3%.

Tsik-ek (ts, n=134): detection 92.5%, assignment 96.0%. Twitter/trill (tr, n=6): detection 66.7%, assignment 100%.

Overlap did not degrade performance in this dataset: for frames labeled as overlapping (n=211), detection and assignment were both ∼100%. Non-overlapping events (n=308) showed detection 94.2% and assignment 95.9%. This pattern suggests robust behavior in multi-caller scenes.

#### Controlled playback for ground truth

In controlled playback, all calls were detected by the camera in both layouts (2-speaker: 100%, 95% CI ≈ 96.0–100%; 4-speaker: 100%, 95% CI ≈ 98.8–100%). Assignment accuracy (correctly attributing a detected call to the emitting speaker) was high in the 2-speaker condition (97.6%; 95% CI ≈ 94.0–99.2%) and lower in the 4-speaker condition (76.3%; 95% CI ≈ 70.9–81.0%). Position-wise patterns were consistent: in the 2-speaker layout, accuracy was ≥95% across front-close, front-spaced, and back-spaced positions (95.0–100%); the hardest case was back-close (90.6%; 95% CI ≈ 84.1–94.6%). In the 4-speaker layout, accuracy remained highest at front-spaced (92.5%; 95% CI ≈ 85.0–96.5%) and front-close (85.0%; 95% CI ≈ 76.9–90.8%), but dropped at back-spaced (66.3%; 95% CI ≈ 56.9–74.6%) and was lowest at back-close (53.8%; 95% CI ≈ 44.1–63.1%). These data indicate that while detection was saturated under all tested geometries, correct source attribution degraded as spatial crowding and rear-plane placement increased, consistent with the beamformer’s reduced separability for nearby or posterior sources.

## Discussion

Understanding who vocalizes, when, and from where is central to linking communication with group decisions and relationships, but continuous, identity-resolved recordings in multi-animal settings remain challenging. Acoustic imaging offers a promising solution by providing spatialized audio without the need to collar every animal. Yet its empirical value rests on four questions: (i) does the sensor is sensitive to the right spectrum at the levels where marmoset calls occur (frequency response & sensitivity)? (ii) how precisely can it localize across the usable field (spatial localization accuracy)? (iii) can it disentangle overlapping sources in space and level (source separation of static, simultaneous emitters)? and (iv) does performance hold when sources move and when real, multi-animal interactions unfold (moving emitters & natural settings)? We addressed these questions sequentially, moving from controlled tests to naturalistic recordings and validating with ground-truth playback, with shared preprocessing and fixed thresholds for reproducibility.

### Frequency response and sensitivity

For typical marmoset call energy (5–10 kHz), the camera behaved close to neutral and was most sensitive (ΔFRD ∼0 dB; lowest detection thresholds), supporting uncorrected mapping and detection in this band (Figure 1). Deviations grew toward 2–5 kHz and 10–20 kHz, consistent with array aperture limits, element roll-off, and SNR. These edges remained useful for detection but are less suited for fine spectral quantification unless compensated (e.g., simple bandwise EQ derived from ΔFRD). Because our ultrasonic reference rolls off <2 kHz, we confined analysis to 2–20 kHz and note that sub-2 kHz work should include a low-frequency reference.

### Spatial location accuracy

Localization accuracy was high and followed a clear spatial gradient: sub-centimeter at the enclosure center and ∼2–3 cm toward edges/corners (Figure 2b). This is the expected footprint of a finite-aperture beamformer: off-axis broadening and stronger early reflections reduce peak sharpness and centroid stability. Call-type effects were modest but consistent with the spectral results. Signals energizing 5–10 kHz with steadier envelopes (e.g., phee) yielded slightly tighter peaks than bursty or spectrally tilted calls (Fig. 2c). Two implementation notes matter for reproducibility: (i) we summarize events via medians across frames to avoid single-frame outliers, and (ii) centroiding from thresholded heatmaps trades off precision and robustness, storing blob area or a peak-to-background metric alongside centroids is a simple way to flag low-confidence frames.

**Figure 2.**
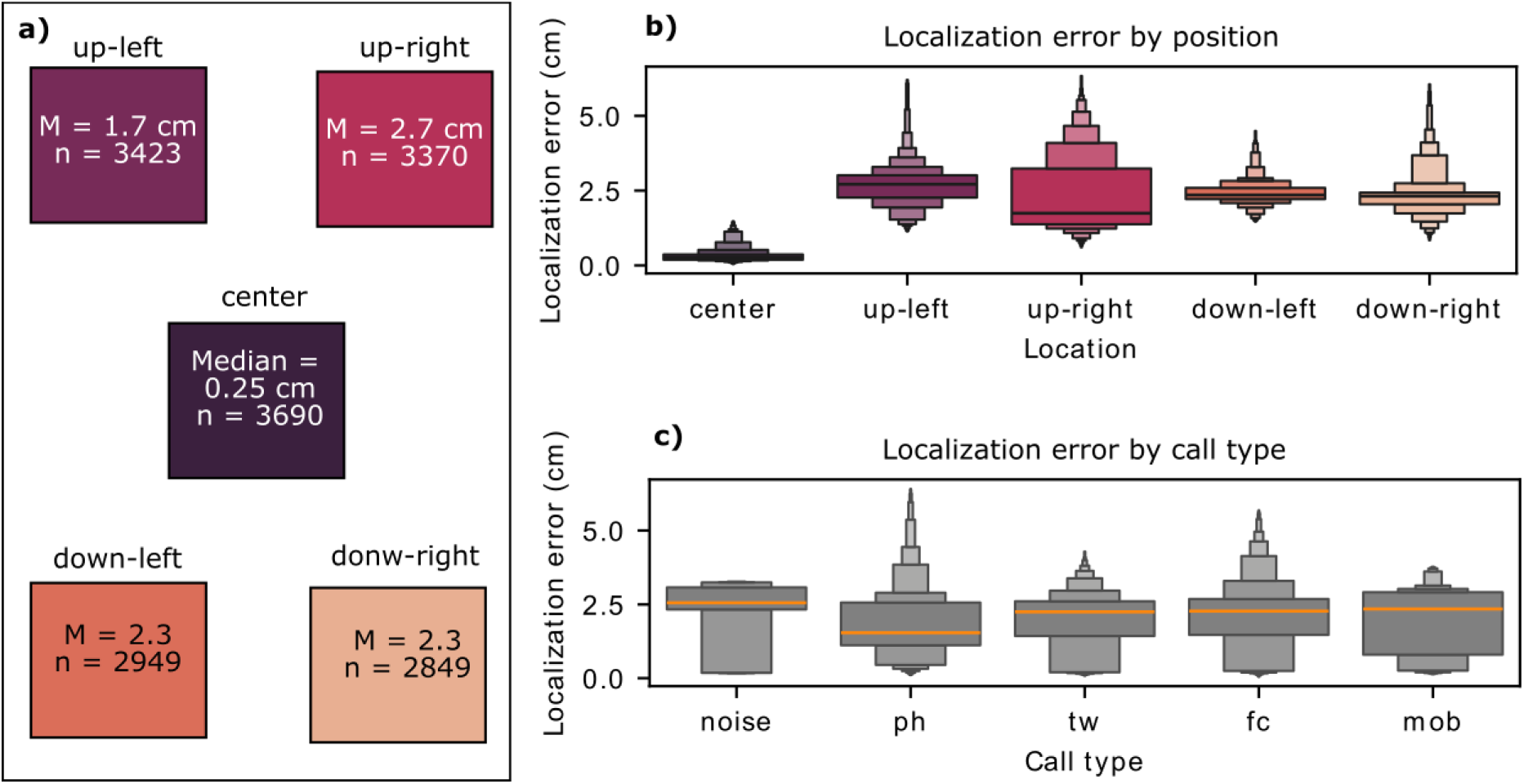
a) Localization by position (“enclosure map”). Median localization error (cm) at the five tested positions (center, up-left, up-right, down-left, down-right). Tiles are color-coded by median error, the numeric median is printed in each tile, with n events shown below. b) Event-level error by position. Distribution of per-event localization errors across positions (boxen plots), box widths reflect data density across quantiles, orange line indicates the median. c) Event-level error by call type (pooled over positions). Distribution of per-event errors for phee, twitter, food, tsik-ek, and band-limited noise (boxen plots, medians overplotted). Errors are Euclidean distances in centimeters from hotspot centroid to the known loudspeaker center. Event-level errors are medians across frames within an event (burst/train).

### Source separation of static, simultaneous emitters

With two simultaneous emitters, separability depended strongly on spacing (Figure 3a-b). At 20 cm, energy collapsed into a single merged hotspot midway between speakers (two “detections” but identical centroids), at 40–60 cm, frames with two distinct peaks were a minority, yet when a side was detected, its localization remained accurate (∼1.6–2.8 cm), at 80 cm, simultaneous two-peak frames were rarer still, but detected centroids were precise. Minute inter-source level differences (≈0.2–1.0 dB) biased the spatial map toward the louder source, often suppressing the weaker lobe below the visualization/detection threshold (Fig. 3d–f). In short, co-representation per frame is fragile, per-source localization when present, is reliable.

**Figure 3.**
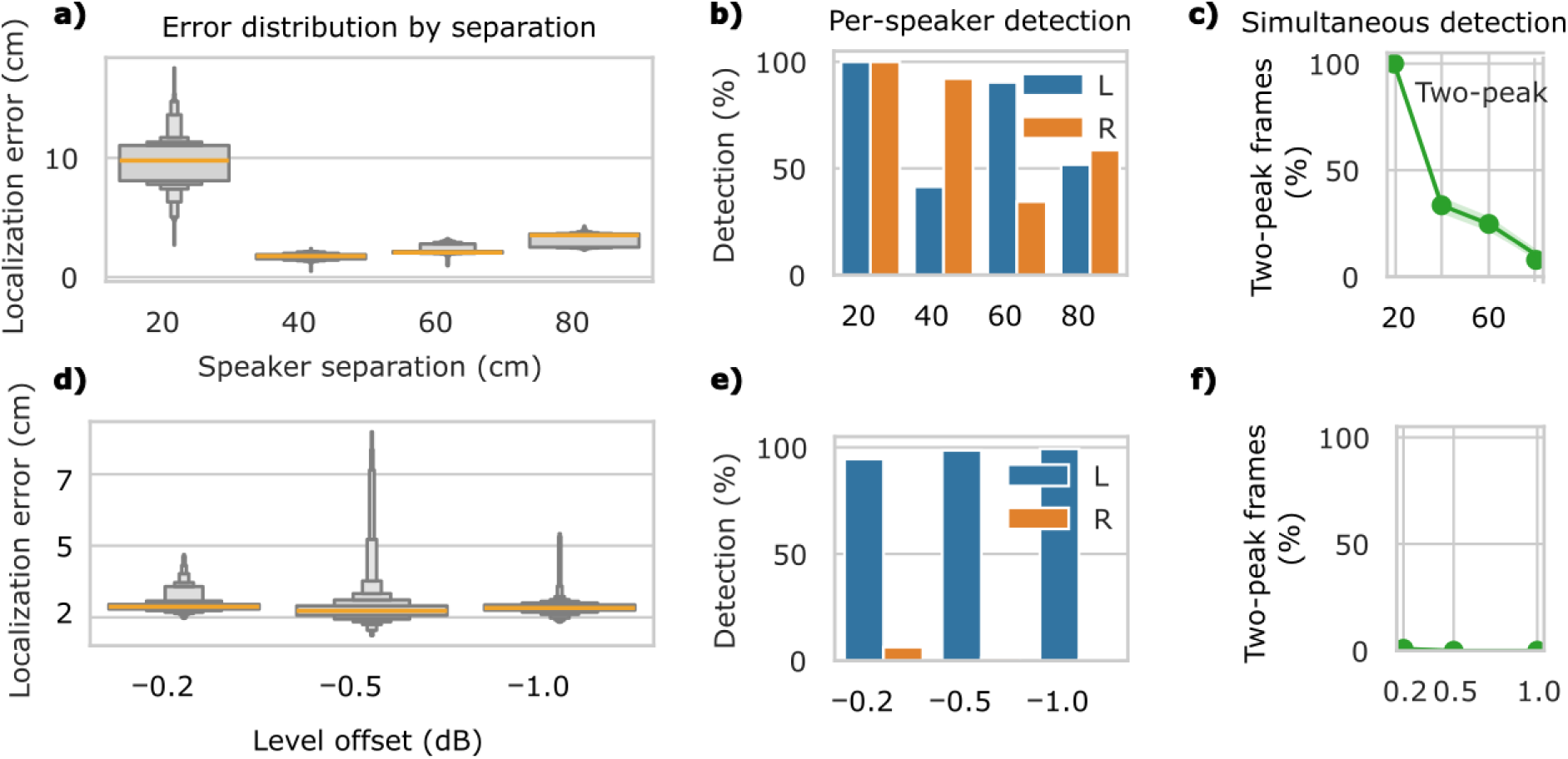
a) Localization error by separation. Box plots show the distribution of centroid-to-speaker distances (cm) for frames where a hotspot was detected. b) Per-speaker detection versus separation. Bars show the percentage of frames with a hotspot detected for the left (blue) or right (orange) speaker. c) Two-peak detection versus speaker separation. Each point is the percentage of frames in which both speakers were detected; shaded bands show Wilson 95% CIs. d) Localization error versus inter-speaker level offset. Box plots summarize per-frame errors across offsets of the test speaker relative to 70 dB SPL. e) Per-speaker detection versus level offset. Bars show the percentage of frames with a detected hotspot for each speaker. f) Two-peak detection versus level offset. Points give the percentage of frames detected by both speakers. Shaded bands are Wilson 95% CIs. Detection is defined framewise as a hotspot within the pre-registered ROI around a speaker, “two-peak” requires detections for both speakers in the same frame. Error is the Euclidean distance (cm) from hotspot centroid to the true speaker position.

Importantly, these limitations did not apply to overlapping naturalistic vocalizations which were correctly detected with very high accuracy. The reason for this is that overlaps there were seldom perfectly synchronous or spectrally identical: tsik-eks are broadband and brief, phees are long and narrow-band, and food calls form variable bouts (Agamaite et al. 2015). This temporal and spectral diversity creates windows in which energy from the target caller is not masked by others, allowing the array to attribute calls with high accuracy. In addition, and very important to highlight is that for the naturalistic testing, we revisited the native software to manually emphasize every single source (frequency span/dynamic range, “silence points”), which further clarified ambiguous scenes.

### Dynamic source separation of moving emitters

When the emitter moves across the field, frame-level localization remains stable at the magnitudes and speeds tested (Figure 4). This complements the static findings: the same beamforming and image-space heuristics that support separation also generalize to motion. Importantly, for marmosets this enables call-aligned trajectories and turn-taking analyses. The main caveats are speed limits (15 FPS cadence), planar motion, and the use of broadband noise. Future work could stress-test higher velocities, depth changes, and natural calls.

**Figure 4.**
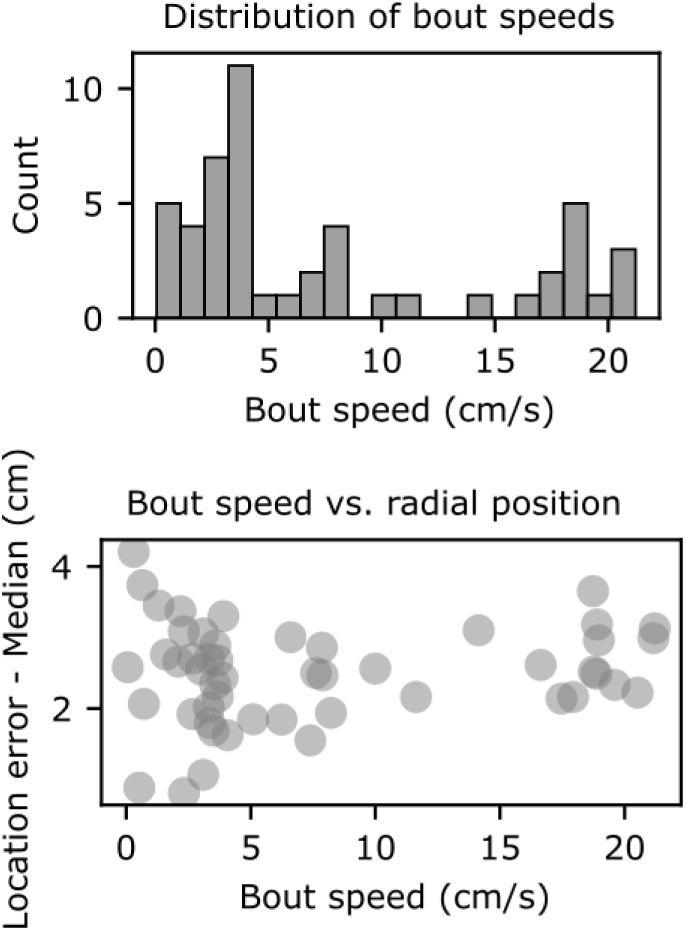
a) Distribution of bout speeds. Histogram shows the speed of each movement bout (cm/s), computed as straight-line displacement of the hotspot centroid between the first and last frame of the bout divided by bout duration. b) Bout speed versus location error. Each point is one bout, x-axis is speed (cm/s) and y-axis is the median distance of the hotspot centroid to the speaker’s geometric center across frames in that bout (cm).

### Real-world marmoset vocal dynamics

In a 15-min family recording (N=519 calls), detection was high (≈96.5%) and, conditional on detection, assignment to the correct caller was ≈97.6%, despite visual clutter, moving animals, and frequent overlap (Fig. 5b-d). Performance varied by call type in line with the earlier sections (food/phee highest; tsik-ek lower), while overlap labels in this dataset did not degrade rates. Ground-truth playback provided a controlled comparison: detection saturated (∼100%) in both 2- and 4-speaker layouts, but assignment decreased as sources crowded the rear plane and increased in number, high with two speakers (≈98%) and lower with four (≈54–92% across layouts, Figure 5e–f). Together, these data indicate that scene geometry and crowding dominate failures in natural scenes. The higher event rate in the 4-speaker playback (≈3 × natural) likely compounded co-activity and assignment difficulty. Note that, unlike the bench tests, ground-truth and natural sessions allowed for recurrent offline adjustment in the native software (narrowing frequency span, tightening dynamic range, setting a “silence point,” or updating array distance) to clarify ambiguous scenes. This accounts for the near-ceiling detection and represents an operator-assisted ceiling, whereas the bench figures quantify fixed-parameter baseline performance.

**Figure. 5.**
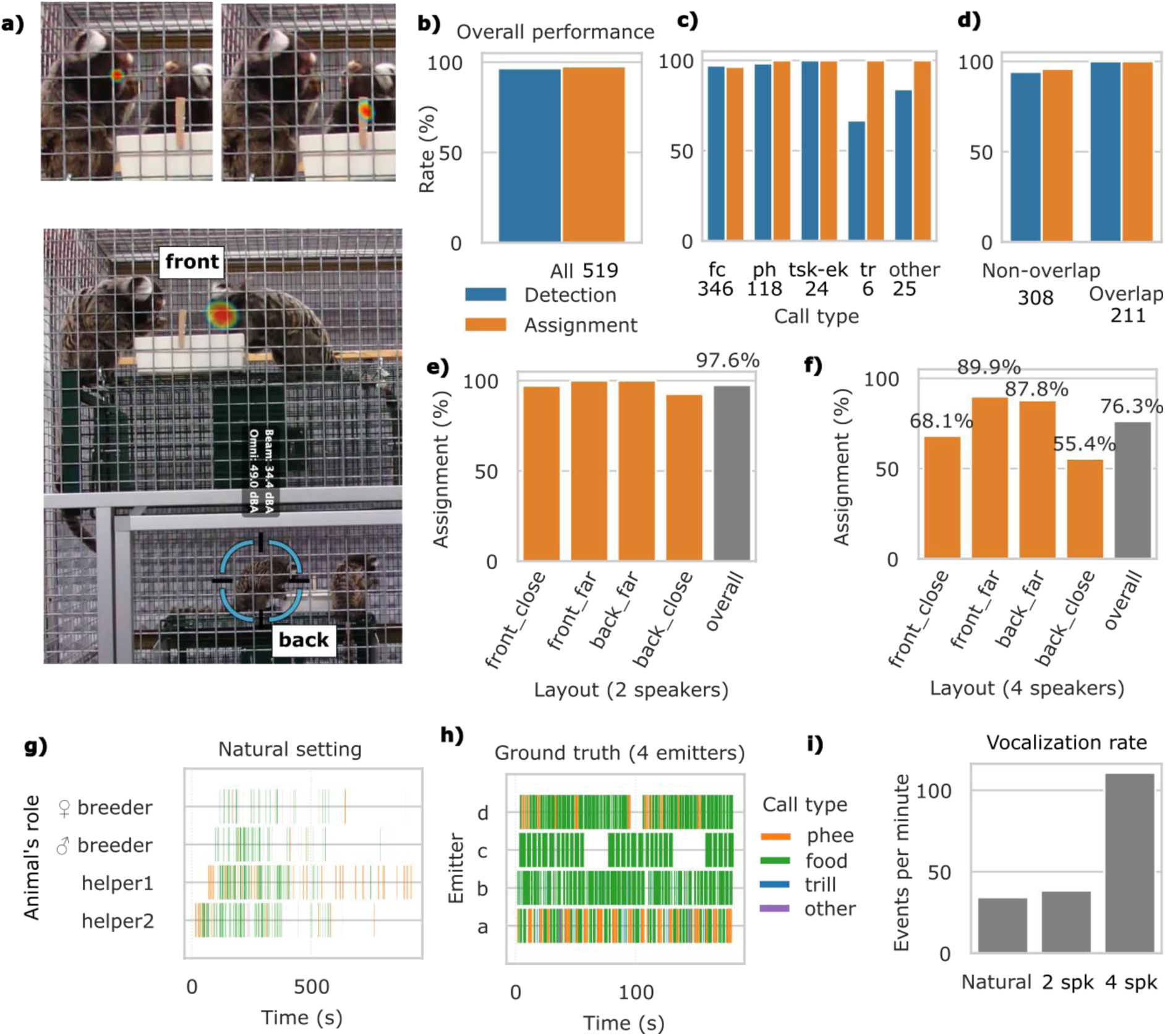
a) Top: Two example frames at different times showing clear hotspots on two consecutive callers. Bottom: layout of the two feeding stations, matching the speaker positions used later in the ground-truth tests. b) Overall performance (natural setting). Bars show detection of camera hotspots for annotated calls and caller assignment given detection. c) By call type. Detection and assignment rates per call (x-axis labels include n). d) Overlap vs non-overlap. Detection and assignment in overlapping vs non-overlapping events. e) Ground truth—two speakers. Assignment accuracy by layout: with an overall detection of 100% across layouts. f) Ground truth—four speakers. Assignment accuracy by layout. g) Natural-setting timeline. Gantt plot of calls over time by animal, colored by call type, illustrating caller alternation, bouts, and overlaps. h) Ground-truth (four emitters) timeline. Gantt plot of the four-speaker playback, showing controlled, labeled call streams per “animal ID”. i) Event rates. Bar plot of vocalization rates (events/min) for natural and ground-truth datasets, providing context for scene density during performance estimates.

Practically, this suggests clear operating guidance. For studies prioritizing whether/when a call occurs (detection), the camera is robust, so automated segmentation plus the heat-map stream yields reliable call counts and timing. For studies prioritizing who called (assignment), two approaches can help in combination. First, shape the scene by positioning the array to favor frontal coverage, maintain ≥20 cm spacing between likely emitters, and place perches/feeding stations where separability is best (front, mid-cage). Second, when crowding is unavoidable, use offline adjudication pass in the native software for ambiguous events by narrowing the frequency span to the call’s band to reduce clutter, tighten the dynamic range so peaks stand out, match the array’s distance to source, and when one source dominates, apply a silence point to momentarily suppress that hotspot so the quitter caller becomes visible in the same frame. These quick, reversible tweaks don’t alter the audio and, in our naturalistic and ground-truth analyses, were often enough to turn uncertain assignments into confident ones.

These results also frame the limits and next steps. First, detection and assignment to specific callers still require manual human-guided work, which is a labor-intensive activity. Integrating multi-animal tracking (pose/ID) and annotation will not just solve this problem but will allow to attach position and interaction state to each call, reducing ambiguity and enabling richer ethology measurements. Second, we validated a four-animal family in a single room. How performance scales to denser groups or neighboring cages (off-camera callers) are unknown. Stress-tests in larger colonies and multi-enclosure rooms, ideally with a second array for triangulation and 3-D localization, should address rear-plane confusions and cross-pen interference. Finally, openness matters. Several of the most useful tools used during testing live behind vendor interfaces. Open programmatic access to key data and controls like intensity coordinates, frequency span, dynamic range, and silence point would let labs reproduce, compare, and extend these analyses at scale.

## Conclusion

Across controlled and natural tests, the acoustic camera performs optimally in the spectral band that carries most marmoset vocal energy, yielding reliable detection and localization. It is most neutral and sensitive in 5–10 kHz, localizes sources with sub-cm accuracy at the enclosure center and ∼2–3 cm near the periphery, and returns precise centroids for detected peaks under concurrency, motion, and real social exchanges. The primary failure mode is not detection but separation of perfectly simultaneous sounds, particularly for closely spaced or rear-plane sources and under slight level asymmetries. Ground-truth playback shows the ceiling is high (near-perfect detection, high assignment with two sources), quantifying the gap attributable to real-world clutter. The next steps are straightforward: scene-aware array placement and lightweight deblending/temporal aggregation for merged hotspots. With these, acoustic imaging becomes a practical tool for continuous, identity-resolved communication in freely behaving primates.

## Funding acknowledgments

This project has received funding from, the European Research Council (ERC) under the European Union’s Horizon 2020 research and innovation programme grant agreement No 101001295 and NCCR: NCCR Evolving Language, Swiss National Science Foundation Agreement no. 51NF40_180888.

